# Adaptive divergence in the bovine genome

**DOI:** 10.1101/022764

**Authors:** William Barendse, Sean McWilliam, Rowan J. Bunch, Blair E. Harrison

## Abstract

Cattle diverged during the Pleistocene into two subspecies, one in temperate and one in tropical environments. Here we have used next generation sequencing of the indicine subspecies of cattle and compared it to the taurine subspecies. Although 23.8 million single nucleotide polymorphisms (SNP) were found, the number of fixed amino acid substitutions between the taurine and indicine subspecies was low and consistent with the Haldane predictions for adaptive selection rather than with Neutral Theory. We noted 33 regions of enhanced divergence of nonsynonymous SNP between the subspecies, which included an increased rate of deleterious variants. Signals of positive selection were found for genes associated with immunity, including the Bovine Major Histocompatibility Complex, which also showed an increased rate of deleterious amino acid variants. The genes important in sensing the environment, especially the olfactory system, showed a network wide signal of positive selection.

## Introduction

Cattle are a rare example of adaptive divergence into temperate and tropical environments in which the subspecies show hybrid vigour on being crossed. Genome wide analyses of evolution in cattle have focussed on the expansion of gene families or on evidence of selection signatures using population genetic evidence based on genotype or haplotype frequencies, mainly in the taurine subspecies. The initial sequencing of cattle found gene families that were expanded in cattle lineages, identifying genes associated with the immune system in particular that were increased in number compared to other mammalian species (Elsik et al. 2009). In addition, analyses of genotype or haplotype frequencies have been used to identify signatures of selection and in some cases have linked these to the known locations of quantitative trait loci (Barendse et al. 2009; Flori et al. 2009; Gibbs et al. 2009; Hayes et al. 2009). Analyses of nonsynonymous and synonymous DNA variants have been performed for a limited number of amplicons for genes expected to be involved in animal production (MacEachern et al. 2009b).

The description of the genomic variation and structure of the *indicus* subspecies lags behind that of the *taurus* subspecies of cattle. The taurine subspecies is of temperate origin while the indicine subspecies is of tropical origin. Individuals of several of the breeds of taurine cattle, including Hereford, Fleckvieh, Holstein, Black Angus, and rare Japanese breeds have been sequenced (Eck et al. 2009; Elsik et al. 2009; Kawahara-Miki et al. 2011; Stothard et al. 2011; Larkin et al. 2012; Tsuda et al. 2013). There has been targeted resequencing of hundreds of individuals to investigate breed diversity for the Holstein, Fleckvieh and other breeds (Jansen et al. 2013; Daetwyler et al. 2014). These genome sequencing studies have documented variation, not only at the single nucleotide polymorphism (SNP) level, but also structural variation including copy number variation (CNV) and major structural polymorphisms (Bickhart et al. 2012). For the indicine subspecies one Brahman, one Nelore, and a handful of Gir animals have been sequenced (Barris et al. 2012; Canavez et al. 2012; Liao et al. 2013) and there is currently no published *de novo* assembly of an indicine genome.

To further our understanding of the indicine subspecies, we have resequenced 32 individuals of the Brahman breed in Australia. Animals were chosen to be as unrelated as possible and were taken to represent a wide sample of the breed in Australia, which originated in the 19^th^ century. These individuals were compared to one another and to sequences from the taurine subspecies. Our aim was to describe the genome wide patterns in nonsynonymous and synonymous mutations to obtain evidence of adaptive selection and divergence between the subspecies.

## Results

We identified 23.8 million single nucleotide polymorphisms (SNPs) different to the Hereford reference sequence across the 32 bulls with an average yield of 5.9 million SNP per individual (Table S3) after collecting 670 Gb of genomic sequence. Of these, 0.62% were coding sequence SNP (cSNP) and 0.25% were nonsynonymous SNP (nsSNP) (Table 1). When normalized for the number of SNP identified in each bull, the proportion of SNP of each type was relatively similar between individuals. However, there was seven times greater variation in the proportion of nsSNP between individuals than for intergenic SNP (Table 1). The degree of similarity between animals and amounts of sequence obtained per individual are contained in the Supplementary Information and Tables S1 and S2. The FASTQ file for each bull was submitted to the 1000 Bulls database (1000Bulls.org) and the VCF file for each bull was submitted to dbSNP under the handle WBARENDSE (BRA2 – SNP and BRA2 – indel) and BioSample object IDs SAMN03166172 to SAMN03166203.

**Table 1.** SNP and indel variability in the sample of Brahman bulls

| **Type** | **Mean** | **SD** | **MIN** | **MAX** | **CV** |
| --- | --- | --- | --- | --- | --- |
| **SNP** |  |  |  |  |  |
| **Intergenic/All** | 0.6628 | 0.0074 | 0.6430 | 0.6789 | 0.011 |
| **NS/All** | 0.0025 | 0.0002 | 0.0022 | 0.0029 | 0.077 |
| **INDELS** |  |  |  |  |  |
| **Intergenic/All** | 0.6498 | 0.0064 | 0.6309 | 0.6610 | 0.010 |
| **NS/All** | 0.00028 | $3.31 \times 10^{-5}$ | 0.00020 | 0.00037 | 0.119 |

The sequencing yielded on average approximately 560,000 small indels per individual (Table S3), of which 0.028% were nonsynonymous coding indels, which was approximately 10 times less than nsSNP (Table 1). The proportion of intergenic indels per individual and the variability between individuals was similar to that found for SNPs. Furthermore, the variability for nonsynonymous indels as a proportion of the total was greater than the corresponding variability for nsSNPs, indicating that individuals differed more substantially from each other in coding sequence indels than in cSNPs. Indels in coding sequence are more likely than SNPs to represent mutations that have strong effects on protein structure and processing.

To characterise the SNP discovery distribution, we plotted the number of Brahman bulls that shared one or more SNP that are different to the cattle reference sequence. Of the 23.8 million SNP, 6.33 million (26.5%) showed either one or two alleles but occurred only in a single animal (Fig 1A). At the other extreme, 12,231 (0.05%) of the SNP were common to at least 27 of the sires, representing SNP that were at high frequency in Brahman animals compared to the Hereford reference sequence. The distribution in Fig 1A appears to reach an asymptote. To determine the decay of the curve we plotted the log number of SNP by log number of bulls (Fig 1B) and found that it was not linear across the entire range, showing that the asymptote decays more quickly than expected. This plot shows a greater deficit of numbers of SNP common to all sires than would be expected given the initial decline in sharing of alleles. To determine whether this might be due to low levels of sequence coverage, the data from the 21 bulls with the most sequence were plotted (Fig S2). This showed much greater linearity of the log-log plot across the range, and in combination with the plot of the rate of change in SNPs shared between bulls (Fig S3) suggests that the accelerated decay is partly due to low genome coverage for some bulls. This lack of similarity would also have a contribution from introgression of some taurine alleles into Brahman cattle (Sanders 1980), which would reduce the number of fixed substitutions between taurine and indicine animals. Extrapolation of the log-log plot in its linear phase suggests approximately 10^5^ fixed differences between taurine and indicine cattle across the genome. The low level of fixed differences between taurine and indicine cattle is unlikely to be due to lack of discovery of SNPs, because not only were 23.8 million SNPs identified, the method of comparison is insensitive to low copy number coverage of a particular polymorphic site, and most importantly, each additional bull added progressively fewer new SNPs and an asymptote was reached. Since it is the rare SNPs that are discovered last, it is unlikely that there is a large number of fixed SNP differences between taurine and indicine breeds that is yet to be discovered.

**Figure 1.**
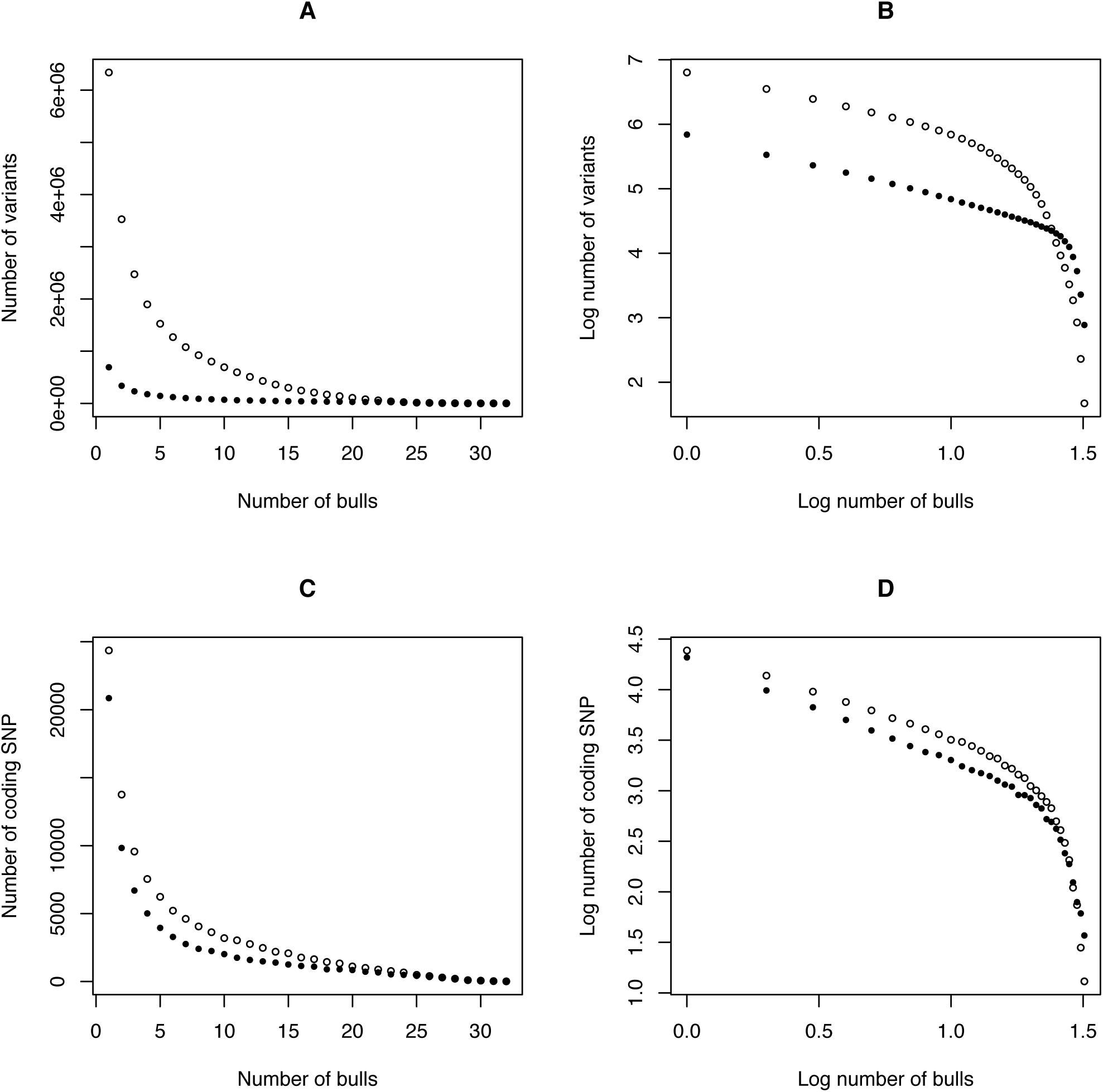
The number of bulls in which a variant is found. **A.** SNP are represented by open circles, small indels by filled circles. **B.** A log-log plot of the data in 1A. **C.** Synonymous substitutions are represented by open circles, nonsynonymous substitutions by filled circles. **D.** A log-log plot of the data in 1C.

Surprisingly, there were more small indels common to all bulls than SNPs common to all bulls (Fig 1B), even though there were fewer indels in total and fewer indels that were found only in a single bull or common to a small number of bulls. The cross-over point was at 23 bulls. There were more than 10 times as many small indels common to all 32 bulls than shared SNPs. The distribution for indels was linear over a larger range of bulls than SNPs, and the decline in number of common small indels was rapid only at the end of the distribution. However, contrary to this pattern of more fixed small indels than SNP, there were no indels in coding sequence that were common to all 32 bulls, and only 5 such indels that were found in more than 27 of the bulls.

Neutral theory predicts that neutral and near neutral mutations, whether slightly advantageous or deleterious, have similar mean times to fixation or extinction (Kimura and Ohta 1971). Therefore, if most nsSNP are neutral or near neutral they should have effectively the same distribution as synonymous SNPs (sSNP). However, we found that there were substantially fewer nsSNP than sSNP in the middle frequency range than at the extremes of the distribution (Fig 1C & D). The log-log plot (Fig 1D) clearly shows that there are more nsSNPs possessed by all 32 bulls than sSNPs even though there are fewer nsSNPs in total. Furthermore, at the other extreme, of SNPs found in only a single individual, the nsSNPs almost equalled the sSNPs. This suggests that a substantial number of the nsSNPs are not neutral or near neutral in their effects. Extrapolation from the linear part of the log-log plot suggests that the number of fixed nonsynonymous substitutions between taurine and indicine cattle is of the order of 900 to 1,000 (Fig 1D, Fig S2, Fig S3). The total number of nsSNPs, irrespective of the number of bulls in which they occur, was 79,309.

To ascertain whether SNP calling, especially the ratio of nsSNP to sSNP, was affected by the draft status of the bovine genome or stringency of SNP calling, we took several approaches to check for errors including examining the location of copy number variants, transcript length and quality threshold for calling of SNPs (Supplementary Information). None of these showed effects that would materially influence the ratio of nsSNP to sSNP in this data set, although we found CNV to have a higher ratio of nsSNP to sSNP than the rest of the genome. We did find that the number of SNPs was strongly correlated to transcript length (Fig 2A, Fig S1), and most genes with large numbers of SNPs have long coding sequences. Furthermore, the number of SNPs per bp was low except for some genes with very short transcript lengths, especially those in the range 50 to 200 bp (Fig 2B). Importantly, we found 33 regions with striking excesses of nsSNP to sSNP (Fig S4, Table S5), and one of these regions included the Bovine Major Histocompatibility Complex (MHC) (Fig S5).

**Figure 2.**
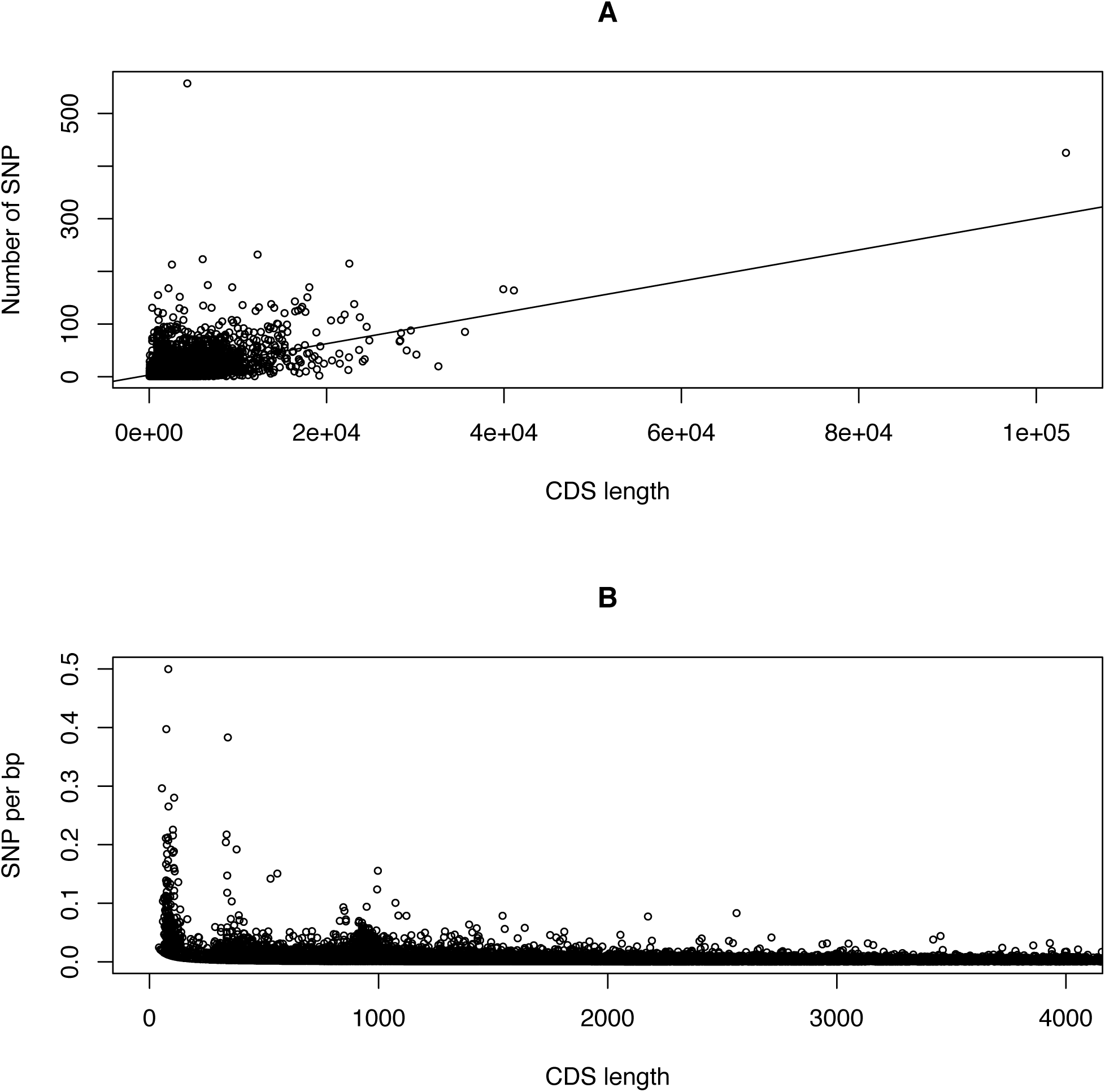
The number of SNP in transcripts of various lengths. **A.** Transcript length in base pairs plotted against number of coding SNP of both types. The line has intercept a=3.29, slope b=0.003, r=0.49 and n=17,881. **B**. Transcript length plotted against SNP per bp, showing excessive SNP per bp is restricted to very short transcripts.

Positive selection is often termed relaxed selection, implying that genes with high mutation rates are more likely to show high levels of nsSNPs. Here we show (Fig 3A) that high numbers of SNP per coding sequence base pair are more likely to be found in values of –0.5log(ω) around zero. A contrast between genes under positive and negative selection (Table 2) shows that although there were significantly (G_adj_ = 736.3, 2df, *P* << 0.00001) more genes under negative selection with low rates of SNP per bp, there were substantial numbers of genes under positive selection that had a low rate of SNP per bp. The average –0.5log(ω) for genes with numbers of SNP per bp ≥ 0.01 (Table S6) was markedly less negative than that for genes with numbers of SNP per bp < 0.01. Although genes with positive selection were more likely to have higher numbers of SNP, there were a large number of genes with evidence of positive selection that had low numbers of SNP per bp and a large number of genes with evidence of negative selection that had a large number of SNP per bp.

**Figure 3.**
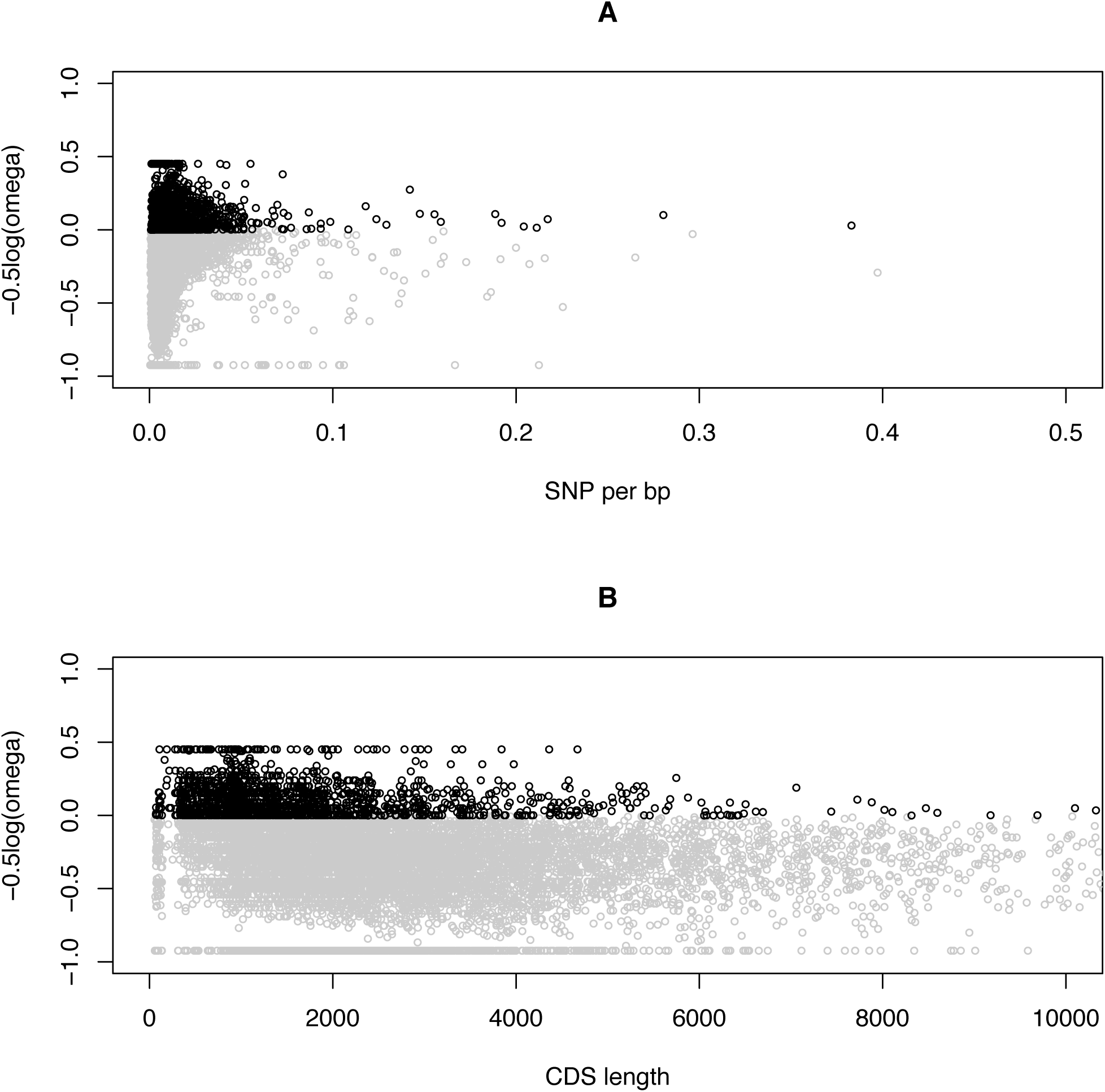
The relationship between positive and negative selection and gene characteristics. **A** Genes with positive selection (black circles) and negative selection (gray circles) plotted against number of SNP per bp in each gene. **B.** Genes with positive selection (black circles) and negative selection (gray circles) plotted against transcript length.

**Table 2.**
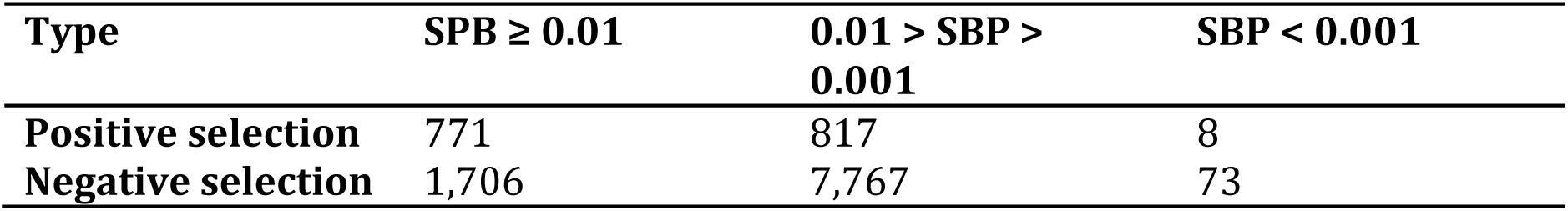
The number of genes under positive and negative selection with low proportions of SNP

| **Type** | **SPB $\geq 0.01$** | **0.01 > SBP > 0.001** | **SBP < 0.001** |
| --- | --- | --- | --- |
| **Positive selection** | 771 | 817 | 8 |
| **Negative selection** | 1,706 | 7,767 | 73 |

Many of the very short genes have large numbers of SNP per bp (Fig 2B) so to examine the impact of numbers of SNP per transcript on estimates of positive selection, the length of transcript was plotted against evidence for positive selection (Fig 3B). Although there is an obvious shift towards shorter transcript lengths for genes with positive selection, this is not driven by transcripts with very high numbers of SNP per bp, such as those with lengths below 500 bp (Fig 3A, Table 3). Removing the transcripts below 500 bp changes the relative length of transcripts under negative selection compared to transcripts under positive selection from 1.74 to 1.65. Shorter transcript length of genes with positive selection appears to be a general pattern in the data and not due to artefacts.

**Table 3.** The average length of genes under positive and negative selection

| **Type** | **N** | **Transcript**<br>**length (bp)** | **S.D.** | **MIN** | **MAX** |
| --- | --- | --- | --- | --- | --- |
| **Positive**<br>**selection** | 1,596 | 1,663.4 | 1,470.1 | 68 | 18,849 |
| **Positive**<br>**selection,**<br>**CDS > 500**<br>**bp** | 1,442 | 1,804.1 | 1,478.2 | 501 | 18,849 |
| **Negative**<br>**selection** | 9,546 | 2,899.9 | 2,769.6 | 54 | 103,346 |
| **Negative**<br>**selection,**<br>**CDS > 500**<br>**bp** | 9,302 | 2,969.1 | 2,772.1 | 501 | 103,346 |

To determine the impact of nsSNP, we characterised the distribution of SIFT predictions. Firstly, fixed nonsynonymous substitutions, and nsSNPs near fixation for opposite alleles, showed SIFT predictions that were very much less damaging than nsSNPs in general. For example, comparing nsSNPs found in 28 or more of the bulls compared to the rest, the average SIFT score = 0.66 S.E.M = 0.017 for those near fixation while the average SIFT score = 0.41 S.E.M. = 0.001 for the rest, yielding t = 37.03, *P* << 0.00001. Using the SIFT codes *deleterious* and *tolerated*, the same comparison yielded an odds ratio of 4.3 for *tolerated* substitutions for those near fixation and G_adj_ = 91.17, 1df, *P* = << 0.00001. Secondly, splitting the data into those transcripts with number of SNP per bp > 0.01 compared to those with SNP per bp ≤ 0.01, irrespective of the number of bulls the SNP was found in, the average SIFT score = 0.40 S.E.M. = 0.002 for SNP per bp > 0.01 while the average SIFT score = 0.42 S.E.M. = 0.003 for SNP per bp ≤ 0.01, yielding t = 5.89, *P* << 0.00001. Using the SIFT codes *deleterious* and *tolerated*, the same comparison yielded an odds ratio of 1.4 for *tolerated* substitutions for those with SNP per bp ≤ 0.01 and G_adj_ = 222.8, 1df, *P* << 0.00001. This comparatively small difference is highly significant due to the very large number of elements. Thirdly, the 33 regions with extremely large numbers of nsSNP compared to sSNP (Fig S4A, Table S5) had genes with 970 deleterious to 3,076 tolerated nsSNP compared to 16,982 deleterious and 58,281 tolerated nsSNP for the rest of the genome, an odds ratio of 1.15 (G_adj_ = 12.95, 1df, *P* = 0.0003). Finally, the distribution of deleterious nsSNPs in transcripts with high SNP per bp was compared to the Poisson distribution and there were substantially more transcripts with either very high numbers or very low numbers of deleterious mutations than expected (Table S7), resulting in a goodness of fit chisquare χ^2_10_^ = 7242.2, *P* << 0.00001. This striking divergence from random was found whether transcripts were segregated on number of tolerated nsSNPs or not (Table S8). A high excess of deleterious nsSNP are consistent with relaxed selection but an excess of zero deleterious nsSNP but high numbers of tolerated nsSNP are implausible under relaxed selection. This suggests other modes of selection, such as balancing selection.

One may predict that genes that are well known, conserved and annotated between species might be subject to more purifying selection. In ENSEMBL, all genes with transcripts have an ENSEMBL name but in some cases these transcripts cannot be linked to known gene identifiers. Of genes with positive selection, 780 had conserved gene names but 816 did not, whereas of genes with negative selection 8,092 had conserved gene names but 1,454 did not. This gives an odds ratio of 5.31 and a G_adj_ = 807.7, 1df, *P* << 0.00001.

To determine whether specific pathways or sets of genes were under selection, genes under positive selection were compared to those under negative selection. Using GO terms and GOrilla (Eden et al. 2009), we found that genes in the immune system and in detection of chemical stimulus involved in sensory perception were significantly represented among the genes under positive selection (Supplementary Information Data File 1). Plots in the region of the Bovine MHC showed elevated numbers of nsSNPs (Fig S5). An examination of the numbers of tolerated and deleterious nsSNP for MHC coding sequences showed much higher than average tolerated (27.75 vs 9.43, t = 5.70 *P* << 0.00001) nsSNPs and slightly higher numbers of deleterious nsSNPs (7.25 vs 3.19, t = 2.66 *P* = 0.008) than other transcripts with large numbers of SNPs per bp.

There was no clear evidence of differences in numbers of genes with excess positive or negative selection at the level of whole chromosomes (Fig 4). One possibility could be the X chromosome, due to initial formation of the Brahman breed using some taurine dams in the 19^th^ century (Sanders 1980), followed by extensive backcrossing using indicine bulls and then the addition of purebred indicine cattle of both sexes to generate an indicine breed. This would predict that the X chromosome would show fewer genes under positive selection, because the comparison is to the taurine reference sequence. On the contrary, we found that the X chromosome had some of the strongest evidence for genes under positive selection. The average –0.5log(ω) across all chromosomes = -0.287 (n=30), S.E.M. 0.007, range -0.354 to -0.163. The value for the X chromosome = -0.163 for n=216 genes, making it the chromosome with the highest level of positive selection, with BTA15 and BTA9 the next highest values (Table S9). It had the 7^th^ lowest number of genes with sufficient cSNP to report values for –0.5log(ω), which was inconsistent with its length and may be a signature from the use of taurine dams.

**Figure 4.**
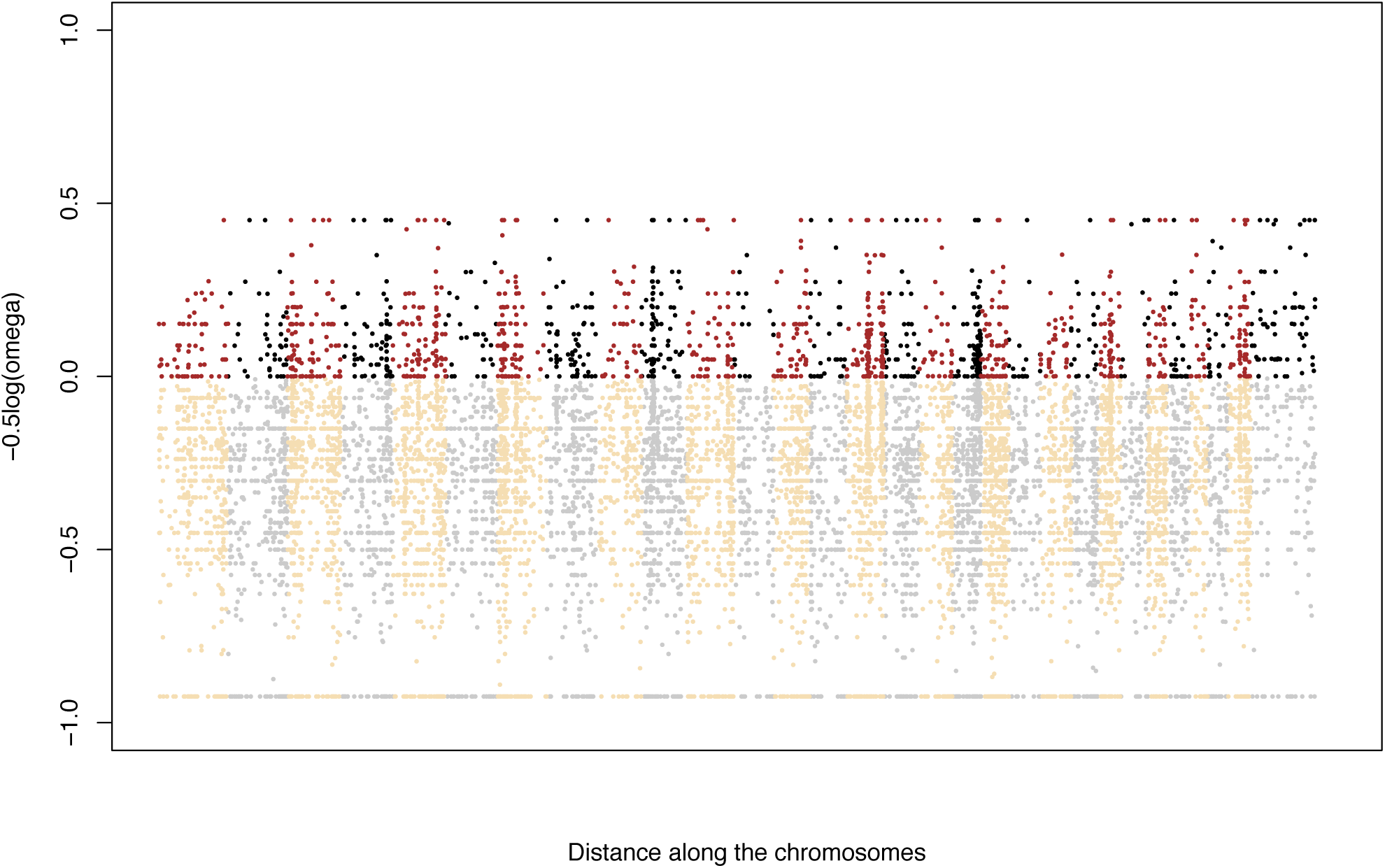
Distribution of –0.5log(ω) in genes across the bovine genome. Positive selection is shown as brown on odd numbered chromosomes and black on even numbered chromosomes. Negative selection is shown as wheat colored on odd numbered chromosomes and gray on even numbered chromosomes.

## Discussion

The analysis of whole genome sequence data from 32 bulls showed that specific classes of genes have evidence for positive selection but surprisingly small numbers of SNPs or indels were fixed between taurine and indicine cattle. We show that shorter genes are more likely to show evidence of positive selection, that signatures of both relaxed and balancing selection were found, and that genes that are under positive selection are approximately 5 times more likely to be poorly annotated than genes under negative selection. We found that genes in the immune system and systems that sense the environment are more likely to show evidence of positive selection. As these subspecies were adapted to temperate versus tropical environments before domestication, these differences likely represent natural selection prior to domestication. Although genes under positive selection were distributed across all chromosomes, there were 33 regions with striking excesses of nsSNPs, which might be regions of focus for divergence between indicine and taurine cattle, and these regions include a slight increase in deleterious nsSNP. Finally, the interindividual variability for intergenic SNPs or indels was low compared to the interindividual variability for nsSNPs and coding sequence indels. As these latter two types of variant are more likely under selection this implies increased variability between individuals for genetic material that is under selective scrutiny. Further research might indicate ways in which this information can be used in prediction of the phenotype.

Surprisingly, only a small proportion of the differences, whether coding or noncoding, SNP or indel, showed fixed differences between Brahman (indicine) cattle and the reference (taurine) sequence. While some of the low number can be attributed to lower levels of genome coverage in some individuals or perhaps to past introgression from taurine germ plasm, if one took as the estimate the lowest number reached in the linear section of the log-log plot this would still only represent 10^5^ fixed differences, much less than 1% of the genetic variants identified in these cattle.

The number of fixed amino acid substitutions between these two subspecies is small and is consistent with adaptive evolution. Using the extrapolations from the log-log plots as noted above, we would expect between 900 and 1,000 fixed nonsynonymous substitutions. The divergence between the taurine and indicine subspecies occurred 0.6 myr to 0.8 myr ago (MacHugh et al. 1997), the effective population size N_e_ prior to domestication is estimated to have ranged between 10,000 to 90,000 (Gibbs et al. 2009; MacEachern et al. 2009a), and the generation interval is approximately 3 years. Using Haldane’s estimate of 1 substitution per 300 generations for horotelic evolution (Haldane 1957), this would yield approximately 600 to 900 fixed substitutions. Kimura estimated the rate of amino acid substitutions based on neutral expectations (Kimura 1968; Kimura and Ohta 1971) and, updating his starting values for values now known for genome sizes of mammals (2.6 Gb) and the size of the combined coding sequence (∼1%), this would result in approximately 1 neutral or near neutral amino acid substitution per 10 years. This would suggest 60,000 to 80,000 nonsynonymous substitutions across the genome. It would require adding together all nsSNPs, including those found only in a single bull, to be consistent with the Kimura prediction. Our results on fixed amino acid substitutions are therefore consistent with the Haldane prediction for genes under standard adaptive selective constraints. Interestingly, these substitutions had four times lower rates of deleterious substitutions than the amino acid substitutions in general. As coding sequences represent around 1% of the genome, extrapolated up to all SNP, the 10^5^ fixed differences between taurine and indicine subspecies are therefore more consistent with polymorphisms under adaptive selection, and would be consistent with large parts of the non-coding sequence showing function, as described by the ENCODE project (Dunham et al. 2012).

Our results show that signals of positive selection are found with shorter genes. If mutations in only part of a gene are under positive selection at any one time, and this is a reasonable expectation given that random amino acid substitutions are more likely to be deleterious, then a measure of overall selection on a gene will be biased to detect negative selection the longer the transcript length of the gene. For very long genes evidence for adaptive selection would be drowned out by the evolutionary constraint on the rest of the molecule.

Some of the evidence for positive selection is consistent with balancing selection although some of it is consistent with relaxed selection. A large number of nsSNP in a transcript could be evidence of relaxed selection, i.e., where genes are not seen by selection on phenotypes and so they accumulate mutations, irrespective of how damaging those might appear to be. Under balancing selection (Wallace 1970), one would expect that many nsSNPs could accumulate on coding sequences that are maintained in polymorphic condition for tens of thousands of generations, but in this case the number of lethal alleles would be constrained. In both cases a high accumulation of nsSNP would lead to an estimate of positive selection even though the nsSNP in some transcripts would have no adaptive significance. Analysis of severity among mutations suggests that there is a 40% increase in deleterious nsSNPS in transcripts with high SNP per bp. Nevertheless, there was both a striking excess of transcripts with high numbers of nsSNPs in some genes where none were identified as deleterious, as well as a striking excess of transcripts in other genes where a large number were identified as deleterious. The former would be consistent with balancing selection and the latter with relaxed selection. One example of balancing selection is the MHC (Bonneaud et al. 2006; Evans and Neff 2009) for which we have evidence of positive selection. In cattle, this complex contains nsSNPs with deleterious effects, which confirms that not all examples of balancing selection will be for alleles with small differences of effect.

## Materials and methods

### Samples

Animal Care and Use Committee approval was not obtained for this study because no new animals were handled in this experiment. The experiment was performed on DNA samples that had been collected previously. DNA samples and duplicates were available for 76 Brahman sires associated with the Cooperative Research Centres for Beef Quality and for Beef Genetic Technologies (Upton et al. 2001; Barwick et al. 2009). These samples were either of industry sires or sires from the Belmont Research Station (Burrow 1998). DNA was extracted from semen or blood (Barendse et al. 2007; Bolormaa et al. 2011). Each sample was genotyped using an Illumina Bovine HD SNP Array (Illumina, Carlsbad, CA) and quality control was performed as previously described (Bolormaa et al. 2011). The genotypes were compared using a genome relationship matrix (Yang et al. 2010) and the 32 least related individuals were chosen for genome sequencing.

### Next generation sequencing

DNA for genome sequencing was checked for quality control using agarose gel electrophoresis, followed by quantification, DNA fragmentation, and size selection using a Nanodrop, a Covaris sonicator, and Qubit-HS DNA kit. DNA for genome sequencing was extracted from whole blood or semen, quantification was by both Nanodrop ND-1000 (Wilmington, DE. USA) and by Qubit dsDNA br Assay, cat no Q32853 by Molecular Probes, supplied by Thermo Scientific, USA. DNA quality was determined by running a subsample of DNA on a 1% 1 X TAE agarose gel to check for DNA smears due to degradation and the 260:280 UV absorbance ratio result on the Nano-drop was required to be between 1.8 and 2.0. Each sample was normalized to 20ng/ul and 55 ul was fragmented to an average insert length of 300 to 400 bp using a Covaris S220 ultra-sonicator (Woburn, MA, USA). DNA libraries for sequencing were prepared from at least 1.0 microgram of genomic DNA of each individual using the Illumina TruSeq DNA Sample Prep Kit v2-Set A kit following the manufacturer’s instructions (Illumina Australia P/L, Scoresby, Vic). A TruSeq PE Cluster Kit v3 was used to generate sequencing clusters using the Illumina cBot on Flow Cell v3. 100 bp paired end reads were obtained using the Illumina TruSeq SBS Kit v3 - HS kit following the manufacturer’s instructions on the SQ module of an Illumina HiScan instrument. Individuals were multiplexed at 8 samples per lane.

### Analysis

Initial quality control consisted of removing sequences that failed chastity filtering using Illumina Casava 1.8. Sequences passing chastity filtering were quality trimmed to remove bases below phred Q20. Using the fastq trimmer module of the Fastx toolkit (http://hannonlab.cshl.edu/fastx_toolkit/commandline.html) sequences less than 50 bp in length following trimming were removed from further analysis. Remaining paired end sequences were aligned to the UMD 3.1 assembly of the cattle genome (Genbank ID GCA_000003055.3), which had been assembled using methods as described (Zimin et al. 2009). The BWA package (Li and Durbin 2009) was used to align the sequence using default values. Alignment included checking that the aligned distance between members of a pair was consistent with the fragment sizes from which libraries were constructed. Sequences were excluded that matched to more than 1 region of the genome. Single nucleotide polymorphisms (SNP) were called using previously described methods and criteria (Barris et al. 2012). The genome sequence variants in the BAM format files were output in VCF format (Li et al. 2009). DNA variation in the VCF files of each individual was annotated to sequence variants using the VEP tool from ENSEMBL (McLaren et al. 2010) and SIFT scores (Kumar et al. 2009) where obtained at the same site. While presence or absence of all the SNP or indels can be ascertained accurately with relatively low sequence coverage it requires 15x coverage to accurately score all the heterozygous bases and 30x coverage to accurately score all the homozygous bases (Bentley et al. 2008; Barris et al. 2012). Therefore the allele frequency data for the majority of scored SNP could be subject to error, whereas identification of the presence of a variant in an animal was of higher certainty. The number of bulls in which a variant was found was therefore calculated noting that this would overestimate the number of fixed differences between taurine and indicine lineages. Numbers and types of each kind of variant were obtained using customized shell and perl scripts. Contingency tables were analysed using G_adj_, the log-likelihood test with the Williams correction (Sokal and Rohlf 1981). Goodness of fit between observed numbers and expectations of the Poisson distribution was calculated (Feller 1968). Pearson correlation coefficients were calculated. Means were compared using unpaired sample t-tests.

The comparison of nonsynonymous and synonymous DNA variants has a long history (Kimura 1968), and an excess of nonsynonymous variants compared to synonymous variants has always been considered as unambiguous evidence for adaptive selection (Kimura 1977; Nielsen and Yang 1998). Most genes, however, do not show higher rates of nonsynonymous than synonymous variants, which has led to many related approaches to identify adaptive selection in genes that are under strong purifying constraint (Kimura 1980; Li et al. 1985; Goldman and Yang 1994; Nielsen and Yang 1998; Suzuki and Gojobori 1999; Yang and Bielawski 2000; Kosakovsky Pond and Frost 2005; Meyer and Wilke 2013). Specifically, these approaches correct for the rate of transition and transversion variants, some use estimated standard errors as part of significance testing while others use the likelihood ratio test. Some models incorporate information on biological differences between variants or information from the branching pattern of evolutionary relationships. It has been shown that the differences in these methods, while methodologically interesting, do not generate much difference in output, suggesting that it does not matter too much which method is used (Kosakovsky Pond and Frost 2005). Here we evaluated the ratio of sSNP to nsSNP (Kimura 1980) where ω = 2k_s_/k_n_ where k_s_ is the value calculated from transitions and transversions for sSNP while k_n_ is the value calculated for nsSNP. Here, genes were included in the comparison if k, the overall statistic over all nucleotides, was at least twice as large as its standard error, which corresponded to genes with at least 4 cSNP. This method has advantages for genome wide analyses. It is conservative. It requires only information on the length of the coding sequence and the number of transitions and transversions for sSNP and for nsSNP. This information is readily available after processing of data into VCF and VEP format and does not require additional realignment of coding sequences. Values below 1 represent an excess of nsSNP. To graph this, however, given the visual disparity between values in the range 0 to 1 versus 1 to ∞, after examining the range of values, ω was plotted as – 0.5log(ω) and values of ω = 0 were truncated at 0.125 while values of ω = ∞ were truncated at 70.6, as these just exceed the values in the distribution of ω over all genes.

This generated a plot where positive selection is represented as values of –0.5log(ω) > 0 and negative selection as values of –0.5log(ω) < 0. GO terms for genes with ω ≤ 1 where compared to GO terms for genes with ω > 1 using GOrilla (Eden et al. 2009), via the website located at http://cbl-gorilla.cs.technion.ac.il/, which implements a Benjamini-Hochberg multiple testing procedure. As there is no specific database for *Bos taurus* or other artiodactyls on this website, even though GO terms are annotated for cattle in ENSEMBL, the comparison was performed for each of the Human, Mouse and Rat databases.

## Acknowledgements

The sequence acquisition was co-funded by the Cooperative Research Centre for Beef Genetic Technologies. We thank M.E. Goddard who supported and encouraged this work. B.P Dalrymple, J.W. Kijas and S.A. Lehnert read and commented on an earlier version of the manuscript.

